# Disruption of the MreB elongasome is overcome by mutations in the TCA cycle

**DOI:** 10.1101/2020.06.18.160713

**Authors:** Brody Barton, Addison Grinnell, Randy M. Morgenstein

## Abstract

The bacterial actin homolog, MreB, is highly conserved among rod-shaped bacteria and essential for growth under normal growth conditions. MreB directs the localization of cell wall synthesis and loss of MreB results in round cells and death. Using the MreB depolymerizing drug, A22, we show that changes to central metabolism through deletion of malate dehydrogenase from the TCA cycle results in cells with an increased tolerance to A22. We hypothesize that deletion of malate dehydrogenase leads to the upregulation of gluconeogenesis resulting in an increase in cell wall precursors. Consistent with this idea, metabolite analysis revealed that *mdh* deletion cells possess elevated levels of several glycolysis/gluconeogenesis compounds and the cell wall precursor, UDP-NAG. In agreement with these results, the increased A22 resistance phenotype can be recapitulated through the addition of glucose to the media. Finally, we show that this increase in antibiotic tolerance is not specific to A22 but also applies to the cell wall-targeting antibiotic, mecillinam.

## 1 Introduction

The bacterial actin homolog, MreB, is a highly conserved and conditionally essential protein necessary for shape and growth in many rod-shaped human pathogens, including *Escherichia coli, Salmonella enterica, Pseudomonas aeruginosa*, and *Vibrio cholerae* (van den Ent, Amos et al. 2001, Alyahya, Alexander et al. 2009). MreB polymers direct the localization of cell wall synthesis, and disruption of MreB, either through the use of antibioitics such as A22 or genetic modifications, causes cells to become misshapen and lyse. A22 binds adjacent to the ATP binding pocket of MreB and alters MreB’s polymerization dynamics (Gitai, Dye et al. 2005, Bean, Flickinger et al. 2009, van den Ent, Izoré et al. 2014, Awuni, Jiang et al. 2016, Awuni and Mu 2019).

Slow growth or overexpression of the cell division genes *ftsZAQ* can overcome the loss of *mreB* (Kruse, Bork-Jensen et al. 2005, Bendezú and de Boer 2008). While overexpression of the cell division genes does not prevent the cell from becoming round upon disruption of MreB, the increased levels of FtsZAQ are thought to help overcome the change in cell diameter, aiding the round cells in dividing. It is currently unknown if over expression or deletion of other genes can also overcome MreB disruption.

There is ample evidence to support a connection between central metabolism and cell shape (Weart, Lee et al. 2007, Elbaz and Ben-Yehuda 2010, Yao, Davis et al. 2012, Hill, Buske et al. 2013, Beaufay, Coppine et al. 2015). In addition, a connection between the TCA cycle and cell size has been shown in *Caulobacter crescentus*. In *C. crescentus*, mutations that result in the accumulation of α-ketoglutarate cause cell shape defects by reducing the synthesis of cell wall precursors (Irnov, Wang et al. 2017).

Similar to cell shape regulation, cell size is coordinated with nutrient availability. Cell size is regulated with growth rate so that cells grown in rich medium are larger than those grown in nutrient poor medium (Schaechter, Maaloe et al. 1958, Sargent 1975, Pierucci 1978). In *Bacillus subtilis*, pyruvate has been shown to help coordinate growth and division through FtsZ-ring assembly (Monahan, Hajduk et al. 2014). Furthermore, in both *E. coli* and *B. subtilis*, the nucleotide sugar UDP-glucose is used as a way to link carbon availability with cell size (Weart, Lee et al. 2007, Chien, Zareh et al. 2012, Hill, Buske et al. 2013).

Because of the well-known connection between cell shape and size and metabolism, specifically the TCA cycle, we screened deletions of genes involved in each enzymatic reaction of the TCA cycle for changes to the minimal inhibitory concentration of A22 (MIC_A22_) in *E. coli*. We found that three gene deletions lead to an increase in the tolerance of cells to A22 and focused our efforts on understanding how disruption of the TCA cycle protein, Mdh, leads to a higher MIC_A22_.

We propose the increase in MIC_A22_ works through the activation of gluconeogenesis leading to an increase in the levels of cell wall precursors. While cell shape changes are not suppressed, we did see an increase to the tolerance of cells to A22 and mecillinam, an inhibitor of the cell wall synthesis enzyme penicillin binding protein 2 (PBP2). Additionally, we found that this higher tolerance for A22 and mecillinam is phenocopied by the addition of glucose to the growth medium. Finally, we found that the cell wall synthesis protein PBP1B is epistatic to Mdh, leading to a reduction of the MIC_A22_ in a PBP1B*mdh* double mutant. These results further support cross talk between metabolism, the cell elongation machinery, and the cell division machinery.

## Results

### Deletions in the TCA cycle result in an increase in the MIC_A22_

It is known that mutations that cause an increase in expression of the cell division genes *ftsZAQ* suppress the growth defects of gene deletions of many cell shape determinants, including MreB (Vinella, Joseleau-Petit et al. 1993, Kruse, Bork-Jensen et al. 2005, Bendezú and de Boer 2008). We wanted to determine if there were gene deletions that result in a cell’s ability to live without the essential bacterial actin homolog, MreB. We used the MreB-depolymerizing drug A22 to mimic the loss of MreB and screened genes from the Keio collection involved in the TCA cycle for mutants that cause an increase in the minimum inhibitory concentration of A22. For quantification purposes we report the concentration of A22 that results in less than half the growth (O.D._600_) compared to cells grown only in LB medium (henceforth referred to as, MIC_A22_) (Iwai, Nagai et al. 2002, Baba, Ara et al. 2006).

We broke each enzymatic step of the TCA cycle by deleting genes for citrate synthase (*gltA*), aconitate hydratase (*acnB*), isocitrate dehydrogenase (*icd*), 2-oxoglutarate dehydrogenase (*sucA*), succinyl-CoA synthetase (*sucC*), succinate:quinone oxidoreductase (*sdhA*), the major anaerobic fumarase (*fumA*), and malate dehydrogenase (*mdh*), (Fig. 1A). Of these, only deletion of *acnB, sucC*, and *mdh* show a higher MIC_A22_ than WT cells (Fig. 1B).

**Figure 1.**
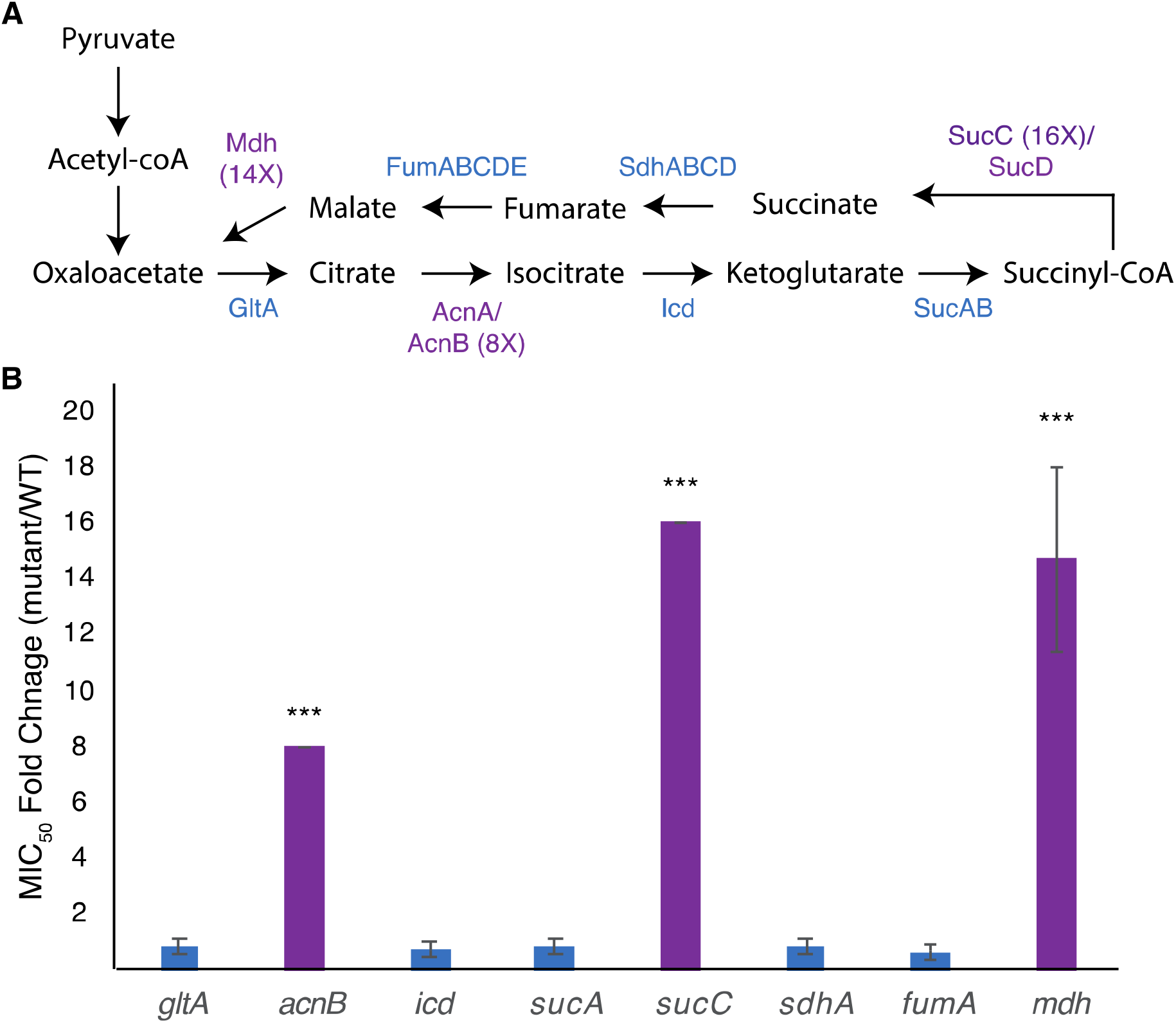
Mutations in the genes of the TCA cycle increase the MIC of A22. A) Model of the TCA cycle. Proteins in purple have higher MIC_A22_ when the associated gene is deleted. B) MIC_A22_ fold change of mutants in the TCA cycle. Only *mdh, acnB*, and *sucC* mutants show an increased MIC_A22_. *** p <0.001 comparing each strain to 1 (no change).

Deletion of *acnB* has been shown to increase antibiotic tolerance by decreasing ATP levels in the cell, but to our knowledge, neither mutations in *mdh* nor *sucC* have been reported to cause a change in antibiotic susceptibility (Rowe, Wagner et al. 2020). In this paper, we focus on understanding the effects of the *mdh* mutation on A22 tolerance. We do not expect the increased tolerance to A22 in the *mdh* mutant to function through ATP synthesis, like *acnB*, because the action of Mdh occurs after the steps in the TCA cycle where reducing power is created.

### Mdh is responsible for the increased tolerance to A22

To ensure that there were not any second site mutations in the Keio *mdh* deletion strain we transfered the mutation into an MG1655 background using P1 transduction. The MG1655 *mdh* mutant retained the increased tolerance to A22 and was used for all subsequent experiments (Table S1). It cannot be ruled out that a second site mutation is genetically linked to *mdh*.

While *mdh* is not in an operon, it is encoded just downstream of the *degQS* operon on the opposite strand. To confirm that the increase in A22 tolerance was from deletion of *mdh* and not disrupted transcriptional termination of *degS*, we complemented *mdh* on an arabinose-inducible plasmid. When expression is induced with 0.2% arabinose the empty vector control has a 6.4 fold increase (p <0.005) in the MIC_A22_ over WT cells compared to a 1.4 fold increase for the complementation strain (p <0.05). When *mdh* is ectopically expressed in the *mdh* deletion cells there is a 4.6 fold decrease (p <0.005) in the MIC compared to the empty vector control, further confirming the ability of ectopically expressed *mdh* to complement the gene deletion. This decrease in the MIC_A22_ when *mdh* is expressed supports the idea that there is not a seconday linked mutation to *mdh* and that deletion of *mdh* results in a higher MIC_A22_. We note that complementation of *mdh* did not fully restore the sensitivity of WT cells to A22, which may be due to either the addition of arabinose or the empty vector as the WT cells induced with the empty vector have a slightly higher MIC than WT cells alone (Table S2).

Slow growth is known to help cells grow without MreB (Bendezú and de Boer 2008). Growth curves of WT and *mdh* cells show that in our test conditions (LB medium) there is no appreciable change in growth rates, indicating that slow growth is not contributing to the increase in MIC_A22_ (Fig. 2A) and that ATP production is not severely affected in this strain. We did however observe that the *mdh* mutant reaches a maximum optical density at a lower point than WT cells. Because the growth rate is the same between WT and *mdh* cells and increased tolerance to A22 is not a universal feature of TCA cycle gene deletions (Fig. 1BC), we hypothesize that the lack of the Mdh enzyme or changes in malate levels (the substrate of Mdh) is the most likely cause of the increased MIC_A22_ seen in the *mdh* mutant; however, it is also possible the lack of oxaloacetate, the product of Mdh is responsible.

**Figure 2.**
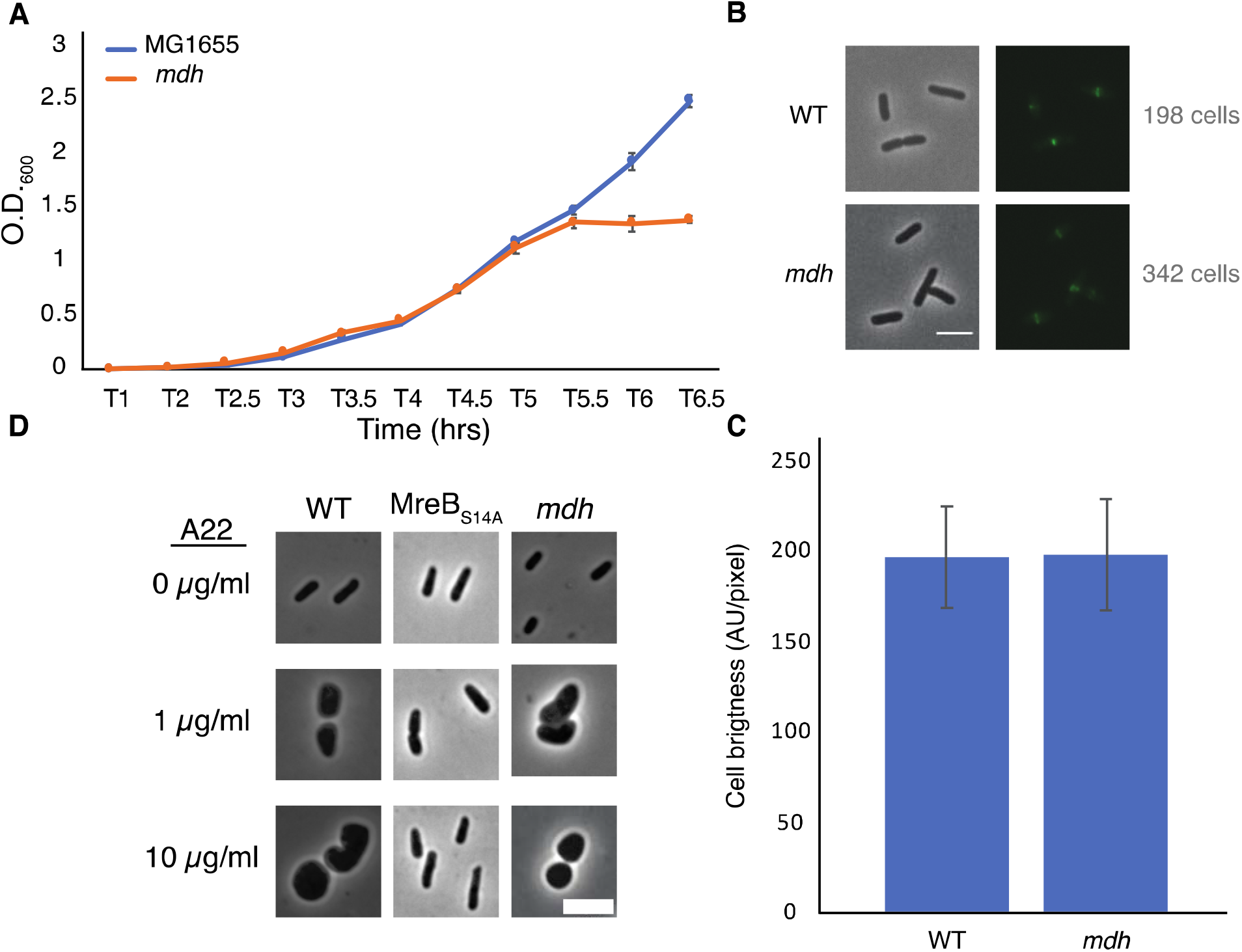
*mdh* deletion cells grow at the same rate as WT cells but lose shape under A22 treatment. A) Representative growth curve of WT and *mdh* deletion cells grown in triplicate. Cells were grown overnight in LB and set to the same O.D. before a 1:1000 dilution was made into fresh LB. Error bars are standard deviation. B) Phase-contrast and GFP images of WT or *mdh* cells with a functional fluorescent FtsZ-GFP fusion. C) Quantification of cell brightness from cells expressing FtsZ-GFP in exponential phase. D) Phase-contrast images of cells grown with different A22 concentrations. Cells were grown for four hours in LB or A22 1 μg/ml. Cells were grown for two hours in LB before 10 μg/ml A22 was added for an additional two hours. Scale bars are 4 μm.

Additionally, it is possible that changes in FtsZ levels are responsible for the increased survival of *mdh* cells. The fact that both the Keio and MG1655 strains display an increase in MIC_A22_ suggests that there is not a second site mutation increasing *ftsZ* levels. It is possible that the metabolic changes caused by the deletion of *mdh* results in changes to *ftsZ* levels. We used a functional native site FtsZ-GFP fusion to measure FtsZ levels in both WT and *mdh* cells. This fusion is the sole copy of FtsZ in the cell. There was no difference in fluorescent intensity between the strains suggesting that FtsZ levels are not affected by deletion of *mdh* (Fig. 2 BC). If FtsZ is overexpressed to a high enough level minicells can form (Belhumeur and Drapeau 1984, Ward and Lutkenhaus 1985). To further support the idea that FtsZ is not upregulated in *mdh* cells, we measured the cell length of non-dividing WT and *mdh* cells and while *mdh* cells are slightly shorter than WT cells, minicells were not produced, suggesting that there are not large changes to FtsZ levels, although the shorter cells do suggest a possible change in divisome activity (Fig. S1A). The lack of minicells can be observed by the lack of small cell outliers (red plus) in the box plot as well as the fact that the whisker representing the most extreme data points is actually lower for the WT cells than the *mdh* mutant, indicating that while the *mdh* mutant produces smaller cells on average the distribution of those cells is also smaller. We also measured the cell length of the *mdh* complementation strain induced with arabinose and found that along with sensitivity to A22 this strain complemented the shorter cell shape phenotype (Fig. 1SB). The loss of *mdh* also leads to a smaller width and cell area, both of which are also complemented when *mdh* is expressed ectopically (Fig. S1C-F).

Cells could also show an increased MIC_A22_ due to a mechanism that prevents A22 from disrupting MreB, as is seen with MreB resistant point mutations. When MreB is disrupted cells become round. To determine if MreB is still being affected by the addition of A22 when *mdh* is deleted, we imaged WT cells, a previously described MreB point mutant resistant to A22 (MreB_S14A_), and *mdh* deletion cells after growth in LB alone, LB with a sublethal concentration of A22 (1 μg/ml) for 4 hours, or LB spiked with 10 μg/ml of A22 for 2 hours after 2 hours of growth (Morgenstein, Bratton et al. 2015). As expected, both A22 treatments cause WT cells to become round, whereas the MreB point mutant remains a rod regardless of A22 treatment. Interestingly, *mdh* cells became round, like WT cells, under both A22 conditions, confirming that MreB is still susceptible to A22 in the *mdh* mutant (Fig. 2D). These data further support the hypothesis that the loss of *mdh* is responsible for the increased MIC_A22_.

### mdh mutants have an altered metabolic profile

We performed metabolomic analysis to identify changes in metabolites in *mdh* mutant cells to determine if these changes might be the cause of the increased MIC_A22_. Both WT and *mdh* cells were grown in LB (without glucose) to exponential phase before metabolite extraction. These experiments were done three independent times and reported results are the ion counts from each of these experiments in order to establish trends in metabolite changes. Only metabolites that show a consistent trend were considered for further analysis. As expected, malate levels were higher in the *mdh* mutant than in WT cells in all three experiments, although the amount varies (Fig. 3A). These increased levels of malate could increase metabolic flux through the alternative malate dehydrogenases MaeA or MaeB, into pyruvate. An increase in pyruvate levels was seen in the *mdh* mutant (Fig. 3B). Pyruvate is involved in many cellular reactions, including glycolysis/gluconeogenesis; therefore, this increased pyruvate could flow into gluconeogenesis to produce sugars (Sauer and Eikmanns 2005). While the levels of glucose-6-phosphate (G6P), the end product of gluconeogenesis, are very low in cells grown in LB and are not seen in one of the trials at all, there was a consistent increase seen in the *mdh* mutant, which could only come from increased gluconeogenesis because the cells were grown in LB medium without any sugar (Fig. 3C). The increased levels of G6P suggest that the elevated levels of malate seen in the *mdh* mutant leads to an increase in gluconeogenesis. G6P can be converted into fructose-6-phosphate which can be used to form N-acetylglucosamine (NAG), while pyruvate can be converted to phosphoenolpyruvate (PEP), another metabolite used in the synthesis of cell wall precursors (Barreteau, Kovač et al. 2008).

**Figure 3.**
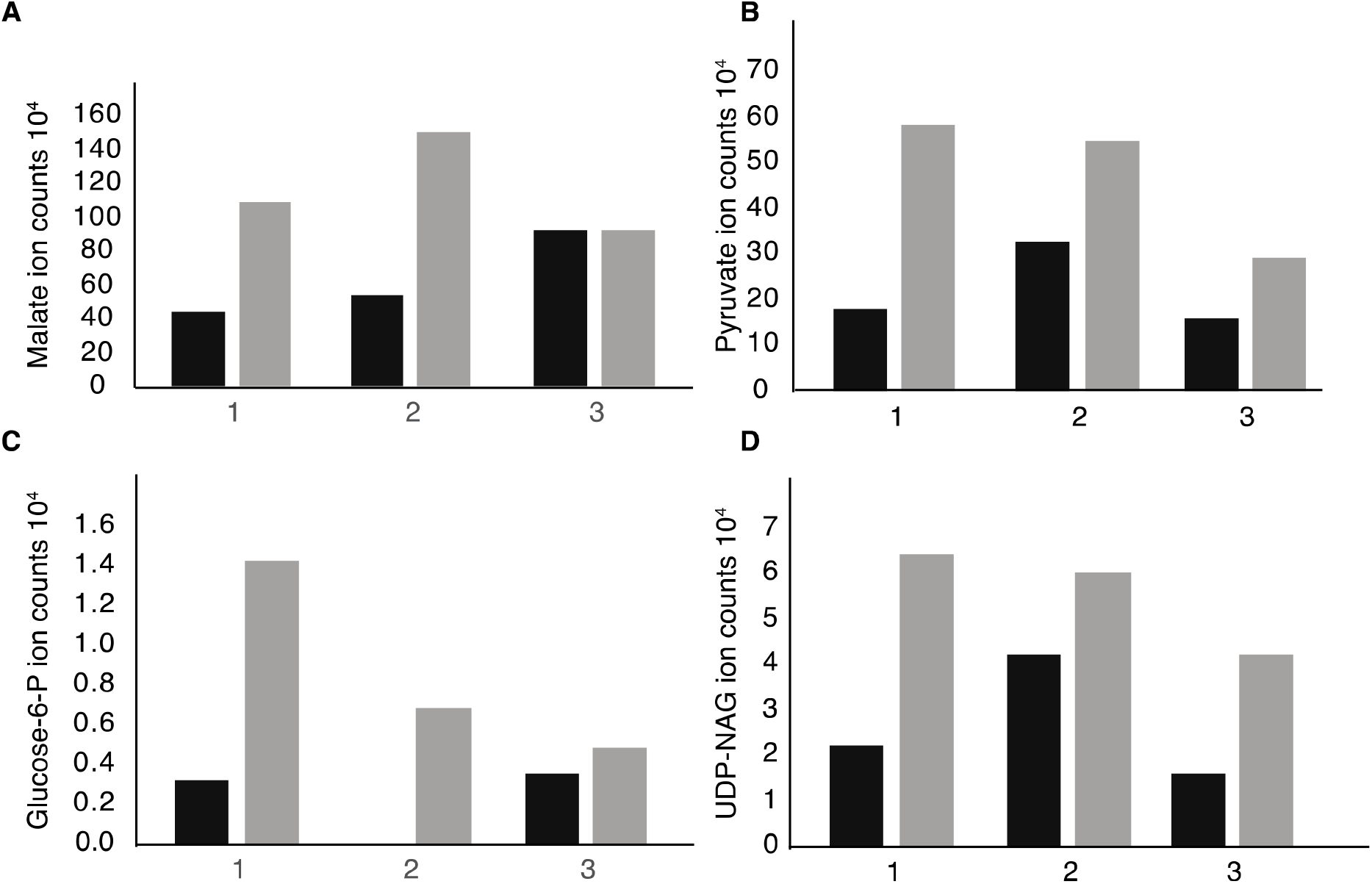
Metabolomics analysis reveals upregulation of gluconeogenesis in *mdh* cells. Cells were grown in LB medium without the addition of sugar on three separate days. The individual ion counts from each day were analyzed to ensure a consistent pattern. A) Ion counts of malate levels in WT and *mdh* cells. B) Ion counts of pyruvate levels in WT and *mdh* cells. C) Ion counts of glucose-6P levels in WT and *mdh* cells. D) Ion counts of UDP-NAG levels in WT and *mdh* cells. Black bars-WT cells, grey bars-*mdh* cells.

Thus, the increased levels of both G6P and pyruvate led us to hypothesize that the *mdh* deletion causes an increase in gluconeogenesis resulting in increased levels of cell wall precursors which may cause an increase in the activity of cell wall synthesis systems not disrupted during A22 treatment (Mengin-Lecreulx, Flouret et al. 1983). To this end, we looked for changes in the level of cell wall precursors in the metabolomics data and saw an increase in UDP-NAG, supporting our hypothesis (Fig. 3D). These results suggest that increasing the amount of cell wall precursors can suppress the lethal effects of MreB disruption.

### The addition of glucose mimics deletion of mdh

Glycolysis and gluconeogenesis are the reverse reactions of each other; thus, because G6P levels increased in *mdh* cells, we reasoned that the addition of glucose to the growth medium would have a similar effect on the MIC_A22_ to deletion of *mdh*: providing additional substrate for the synthesis of cell wall precursors. In *Staphylococcus aureus* it was shown that ~50% of exogenously added glucose ended up as part of the cell wall, supporting the hypothesis that exogenously added glucose can be easily converted into UDP-NAG and incorporated into the cell wall (Komatsuzawa, Fujiwara et al. 2004). Additionally, glucose has been shown to compensate for mutations in aconitase to restore antibiotic susceptibility (Rowe, Wagner et al. 2020).

MIC assays were performed on WT cells grown in LB medium with increasing amounts of glucose or α-methyl-D-glucoside (αMG), a nonhydrolyzable form of glucose (Rogers and Yu 1962, Hernandez-Asensio, Ramirez et al. 1975). While very low levels (<0.125%) of glucose had no effect on the MIC_A22_ in WT cells, even glucose levels as low as 0.125% resulted in a statistically significant increase in the fold change of the MIC_A22_ in cells grown with glucose vs LB alone compared with cells grown with αMG vs LB (Fig. 4A). An increase in the glucose levels results in even higher MIC_A22_ fold changes. The significant increase in MIC_A22_ during glucose vs αMG treatment suggests that the changes are not due to osmotic protection caused by the addition of sugar. These results further support our hypothesis that increasing the levels of cell wall precursors leads to an increase in MIC_A22_.

**Figure 4.**
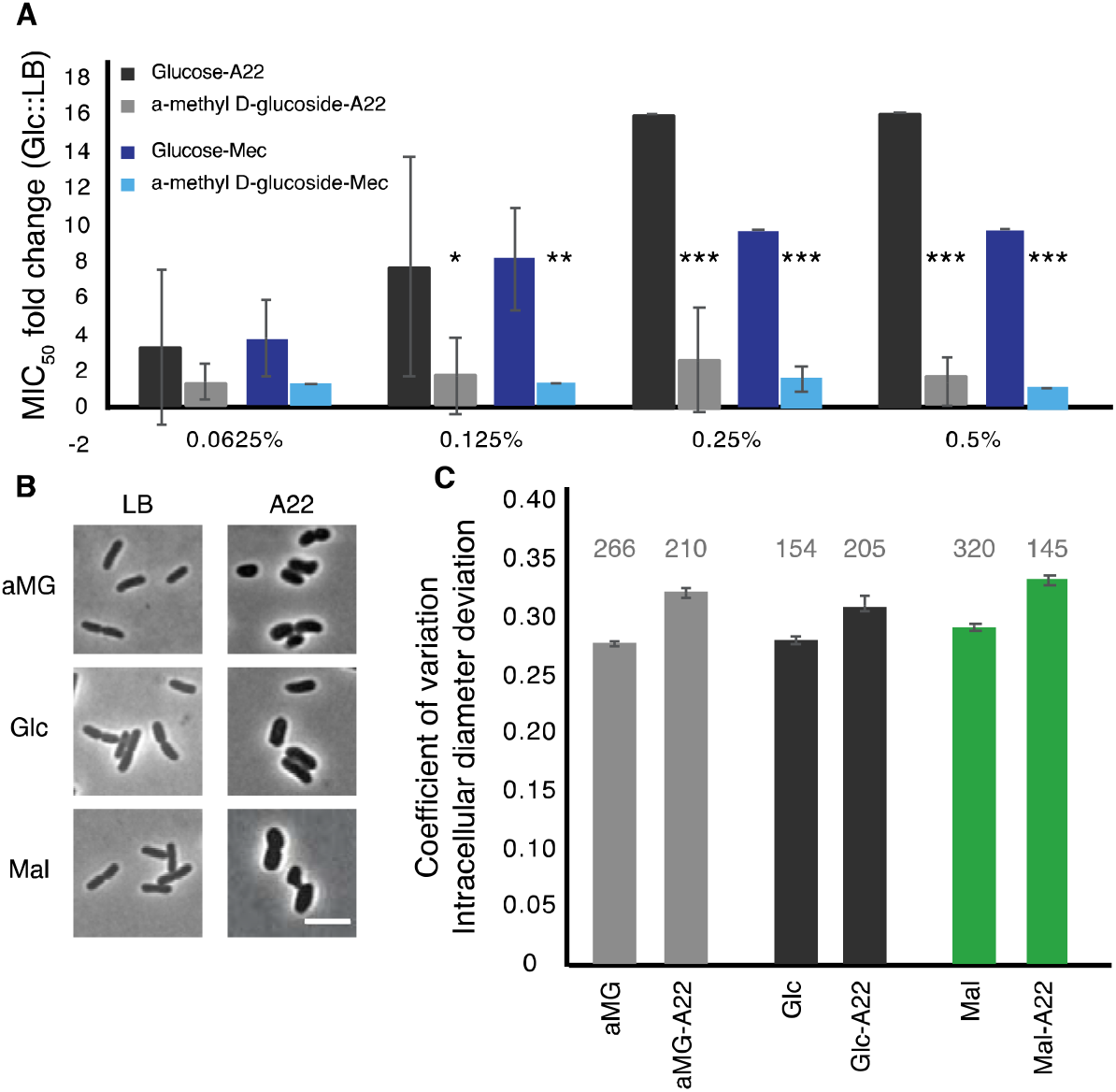
The addition of glucose to the medium phenocopies the antibiotic tolerant phenotypes of the *mdh* deletion. A) WT cells were grown in LB medium supplemented with the indicated amount of either glucose or a non-hydrolizable glucose mimic, α-methyl D-glucoside (αMG). Cells were treated with either A22 (black bars) or mecillinam (blue bars). Error shown is standard deviation from 3 independent experiments. * p<.05, ***p<.001 comparing glucose to αMG for each condition. B-C) Cells were grown with 0.2% αMG, glucose (glc), or malate (mal) for four hours before imaging (LB) or for two hours before a spike of 10 μg/ml A22 for another two hours. B) Images of cells. C) Quantification of how rod-like cells are. Coefficient of variation of the intracellular diameter deviation measures rodness. The higher the number the less rod-like cells appear. A22 causes cells to round up in all conditions but the effect is less when cells are grown in glucose. Number above each bar represents the total number of cells analyzed

Because glucose was able to provide WT cells some protection from A22 we wanted to determine how it affected cell shape. WT cells were grown in LB or A22-spiked media supplemented with 0.2% glucose, αMG, or malate. In all conditions A22 caused a loss of rod shape measured by increases in the intracellular diameter deviation (IDD) (Fig. 4BC) (Morgenstein, Bratton et al. 2015). We observed that the change in IDD was smaller (although still statistically significant) when cells were treated with glucose than either αMG or malate. Using a two-way ANOVA analysis, we are able to show that the addition of glucose has a significant effect on reducing the change in IDD when cells are treated with A22 (p < 0.05).

### Deletion of mdh leads to an increased MIC of mecillinam

The primary role of MreB is to direct the location of cell wall synthesis enzymes. It has been suggested that MreB forms a complex with the SEDS family of cell wall synthesis enzymes (Cho, Wivagg et al. 2016, Meeske, Riley et al. 2016, Leclercq, Derouaux et al. 2017). Specifically, recent work has suggested that MreB interacts with RodA and PBP2 to regulate cell elongation, while PBP1A/1B work independently of MreB (Cho, Wivagg et al. 2016). Because the *mdh* deletion results in cells with a higher MIC_A22_, we hypothesized that cells might also have a higher MIC against other cell wall targeting antibiotics. To that end, we performed MIC assays with other antibiotics that target cell wall synthesis proteins to determine how general the effects of the *mdh* deletion are on the MICs of cell wall-targeting drugs.

We performed MIC assays with cell wall-targeting antibiotics that inhibit either cell elongation (mecillinam, cefsulodin), cell division (cephalexin) or both (ampicillin) by blocking the activity of different PBPs involved in cell wall synthesis. An MIC fold change >1 indicates that the *mdh* deletion causes cells to be more resistant and a fold change <1 indicates that *mdh* cells are more sensitive to the specified antibiotic. The only other cell wall-targeting antibiotic to which the *mdh* deletion shows an increased MIC is mecillinam, which targets the MreB complex partner, PBP2 (Spratt 1977) (Fig. 5A). The fact that the *mdh* deletion strain does not have a higher MIC_cef_ supports the idea that PBP1A/B are not part of the MreB complex because cefsulodin specifically targets PBP1A and PBP1B. Cephalexin targets PBP3 (FtsI) which is part of the division machinery, a different complex than the MreB-elongasome. We did not see an increase in the MIC_ceph_ in the *mdh* deletion, suggesting that an increase in gluconeogenesis cannot overcome cell division inhibition. Ampicillin targets multiple PBPs, with the highest affinity for PBP4, followed by PBP3, and PBP2 (Curtis, Orr et al. 1979, Preston, Wu et al. 1990). Because ampicillin binds to PBP3 tightly and there was no increase in the MIC_ceph_, it stands to reason that deletion of *mdh* would not have an effect on the MIC_amp_. These results suggest that the up-regulation of gluconeogenesis caused by the *mdh* deletion specifically causes an increase in the MIC of drugs that target the MreB elongation synthesis complex (A22 and mecillinam), but not to other cell wall-targeting antibiotics.

**Figure 5.**
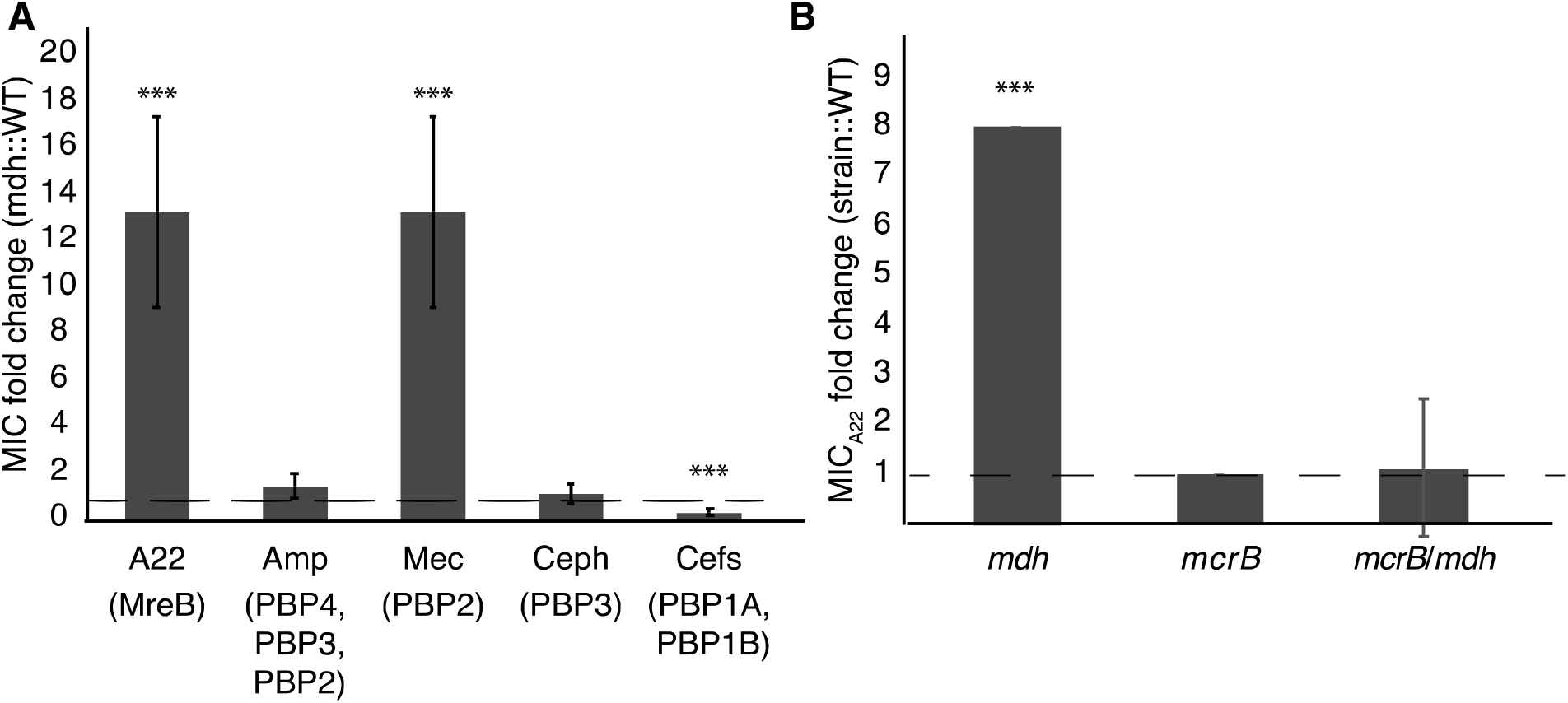
*mdh* mutation provides increased MIC specifically of A22 and mecillinam. A) MIC assays were performed for the indicated antibiotics. Amp- ampicillin, Mec- mecillinam, Ceph- cephalexin, Cef- cefsulodin. A fold change of 1 (dotted line) indicates no change. B) A22 MIC comparing WT cells with the indicated strains. *** p <0.001 comparing each strain to 1.

Inactivation of aconitase (*acnB*) or succinyl-CoA synthetase (*sucC*) lead to an increase in the MIC_A22_ (Fig. 1D); therefore, we tested whether deletions of these genes also lead to an increase in the MIC_mec_. The *acnB* mutation displays an increased MIC_mec_, while deletion of *sucC* does not (Fig. S2). This suggests that the role of SucC in A22 resistance is different f that of Mdh. However, it is unclear if AcnB acts through a general antibiotic resistance mechanism via ATP depletion or through a specific MreB-elongasome mechanism.

### Glucose phenocopies the mdh mutation leading to an increased MIC of mecillinam

The addition of glucose to the media was able to phenocopy the effects of deleting *mdh* on the MIC_A22_ (Fig. 4A). Because we saw a similar increase in the MIC_mec_ as with the MIC_A22_ in the *mdh* deletion, we wanted to test whether the addition of glucose would also lead to an increased MIC_mec_ in WT cells. The addition of glucose leads to a statistically significant increase in the MIC_mec_ but not αMG (Fig. 4). Taken together, the fact that there is only a change in the MIC for antibiotics that target MreB and PBP2 (the major elongasome components), but not other PBPs suggests that an increase in the levels of cell wall precursors, either through an increase in gluconeogenesis or the addition of glucose to the medium, induces a mechanism that specifically targets the MreB elongasome, but not other cell wall synthesis enzymes.

### PBP1B is epistatic to mdh for A22 tolerance

If there are two cell wall synthesis machinery systems at work, it stands to reason that inhibition of one could be compensated by activation of the other. Cefsulodin specifically targets the bifunctional PBP1A and PBP1B proteins which have been suggested to work separately from MreB-PBP2, (Cho, Wivagg et al. 2016, Meeske, Riley et al. 2016). The *mdh* mutation does not increase the MIC_cef_; therefore, to test if either PBP1A or PBP1B can compensate for disrupted MreB we performed MIC assays on *mdh* cells with either *mrcA* (PBP1A) or *mrcB* (PBP1B) deleted. The *mrcB* deletion, either alone or with *mdh*, made cells highly sensitive to A22 (Fig. 5B). Interestingly, while WT cells display reduced growth above 1.25 μg/ml A22, cells lacking PBP1B show no growth above this concentration but did not have increased sensitivity at lower concentrations resulting in the same MIC when measuring 50% of growth of cells versus growth in LB. We hypothesize that the increased levels of cell wall precursors observed in the *mdh* mutant can be used by PBP1B to keep cells alive in the absence of the MreB-PBP2 complex.

## Discussion

While studying novel mechanisms of A22 resistance, we found that loss of Mdh, an enzyme in the TCA cycle, leads to an increase in the MIC_A22_ (Fig. 1), which we hypothesize is through the induction of gluconeogenesis, leading to an increase in cell wall precursors. We propose that this accumulation of cell wall precursors results in increased MICs of both the MreB-targeting drug A22 and the PBP2-targeting drug, mecillinam through the activation of PBP1B. This effect appears to be specific for the MreB cell wall synthesis complex, as we do not see an increase in MIC of antibiotics that target other enzymes in the cell wall synthesis pathway.

Mutations in the TCA cycle have been implicated in resistance to antibiotics. Deletion of *icd*, which encodes isocitrate dehydrogenase, results in the accumulation of toxic intermediates leading to the activation of the AcrAB-TolC efflux pump which can pump nalidixic acid out of the cell (Helling, Janes et al. 2002). A similar mechanism is most likely not at work in the *mdh* mutant, as the cells still become round upon A22 treatment, suggesting that A22 is in the cell at a high enough concentration to disrupt MreB. More recently, it was shown that antibiotic tolerance can be induced by inhibiting ATP production (Conlon, Rowe et al. 2016, Shan, Brown Gandt et al. 2017). One way this could be achieved is by blocking enzymes in the TCA cycle such as aconitase (*acnB*) or succinate dehydrogenase (*sdhA*), which would lower the reducing power available for oxidative respiration (Rowe, Wagner et al. 2020).

### Multiple systems involved in rod shape maintenance

The bacterial cell wall is a macromolecule made up of sugars crosslinked by proteins to provide shape and support to the cell. In rod-shaped bacteria, such as *E. coli, B. subtilis, and C. crescentus*, MreB is thought to form the main protein of the elongasome complex consisting of multiple PBPs, MreBCD, RodA, and RodZ. Work in the above species led to the hypothesis that MreB interacts with the cytoplasmic cell wall synthesis components and helps to direct the localization of the periplasmic-acting enzymes (Figge, Divakaruni et al. 2004, Divakaruni, Loo et al. 2005, Kruse, Bork-Jensen et al. 2005, Divakaruni, Baida et al. 2007, Kawai, Daniel et al. 2009, White, Kitich et al. 2010, Ursell, Nguyen et al. 2014, Morgenstein, Bratton et al. 2015).

Recent work has questioned the model that the MreB elongosome is a large complex composed of both the bifunctional class A PBPs (aPBPs) and monofunctional class B PBPs (bPBPs). The aPBPs contain both transglycosylation and transpeptidation activities, and thus should be functional alone, while the bPBPs, such as PBP2, possess only transpeptidation activity. However, PBP2 is essential in *E. coli*, and in both *B. subtilis* and *E. coli*, polymerization of glycan strands has been shown to proceed without the aPBPs (Cho, Wivagg et al. 2016, Meeske, Riley et al. 2016). If elongation of glycan strands occurs without aPBPs, then there must be another enzyme capable of transglycosylation reactions. RodA has been suggested to be the transglycosylase that works with PBP2 (bPBP) (Cho, Wivagg et al. 2016, Meeske, Riley et al. 2016). These authors also showed that the RodA-PBP2 complex works with the MreB cytoskeleton while the aPBPs act separately from this complex. Our previous work studying MreB dynamics suggested that RodZ helps modulate MreB motion through interactions with RodA/PBP2, further supporting the idea that they are in a complex together (Morgenstein, Bratton et al. 2015). Here we show that antibiotics that target MreB (A22) or PBP2 (mecillinam), but not aPBPs (cefsulodin) are less effective when *mdh* is deleted and cell wall precursor synthesis levels are proposed to be upregulated (Fig. 5A), suggesting that a second PG synthesis system is dominating.

These results reinforce the model that MreB forms a cell wall synthesis complex with PBP2 but not PBP1A/B and suggest that cell wall synthesis deriving from the activity of aPBPs may be upregulated when there is an abundance of precursor molecules in the cell and the MreB elongation machinery is disrupted. Upregulation of an aPBP-specific synthesis mechanism would explain how cells can grow when MreB or PBP2 are inhibited. To test this idea, we made an *mdh* double deletion with either *mrcA* (PBP1A) or *mrcB* (PBP1B). Surprisingly, loss of PBP1B either alone or with *mdh* results in cells more sensitive to A22 (Fig. 5B).

To the best of our knowledge, the rate-limiting step of cell wall synthesis is currently unknown as it is very difficult to measure *in vitro* cell wall synthesis, which is further complicated by the rapid turnover of the PG. Possibilities include substrate availability (precursor synthesis), enzyme (PBPs) kinetics, or enzyme availability. Our results suggest substrate availability is rate limiting, as increasing precursor synthesis provides a mechanism bypass the need for PBP2 or MreB. Future, experiments that reduce substrate levels, through modulation of the synthesis enzymes, or that modulate PBP levels will be needed.

It is possible that the observed increase in UDP-NAG actually results from a decrease in cell wall synthesis. This model would also be consistent with cells becoming more sensitive to the inactivation of PBP1B; if cell wall synthesis is slowed in the *mdh* mutant, loss of another cell wall synthesis enzyme (PBP1B) would be more deleterious than in WT cells. However, it is unclear why the addition of glucose to the medium would result in increased tolerance. How aPBP activity is modulated when other cell wall synthesis mechanisms are inhibited is currently not known but would be an interesting topic for further study. If reduced wall synthesis was happening, one would expect that deletion of PBP1A to also reduce the MIC_A22_ in the *mdh* deletion background, but this was not observed.

While MreB and PBP2 may act in the same complex, inhibition of each protein results in different effects on cell wall synthesis. MreB acts upstream of cell wall synthesis to organize the complex; therefore, A22 treatment breaks the ability of the cell to build an organized cell wall. PBP2 is an enzyme that actually builds the cell wall acting upstream of MreB and A22 inhibition. Inhibition by mecillinam, and other PBP inhibitors, causes a futile cycle where not only is synthesis or crosslinking inhibited but cell wall recycling is increased (Uehara and Park 2008, Cho, Uehara et al. 2014). While most antibiotics that target cell wall synthesis enzymes result in this futile cycle, we only see an increase in resistance to mecillinam, suggesting a specific phenotype for inhibition of the MreB-PBP2 elongasome. We propose that if precursor synthesis is elevated and MreB or PBP2 is inhibited, then the increased activity of PBP1B can break the futile cycle by using the excess precursors.

### Drug efficacy is affected by the metabolic state of cells

Antibiotics resistance normally comes from genetic changes in a cell. However, in addition to genetic changes the efficacy of antibiotics changes with the growth state of cells. Slow growing cells show an increased tolerance to many antibiotics (Tuomanen, Cozens et al. 1986, Pontes and Groisman 2019). When only a subfraction of the population is tolerant to antibiotic exposure, the surviving cells are termed ‘persisters’ (Brauner, Fridman et al. 2016). In addition, changes in metabolic state can affect the efficacy of antibiotics. Previous studies have shown changes to antibiotic sensitivity when mutations are made in the genes involved in the TCA cycle (Helling, Janes et al. 2002, Irnov, Wang et al. 2017, Rowe, Wagner et al. 2020). Mutations in *icd* can activate efflux pumps rendering cells more tolerant to antibiotics that can be targeted by those efflux pumps (Helling, Janes et al. 2002). Additionally, *acnB* mutations are thought to alter antibiotic susceptibility through decreases in cellular ATP levels (Conlon, Rowe et al. 2016).

Here we show that mutations in the metabolic genes *mdh, sucC, and acnB* increase a cell’s tolerance to A22. Both *mdh* and *acnB* mutations also lead to increases in mecillinam tolerance, while *sucC* deletion does not affect mecillinam tolerance (Table S2). This suggests that the changes in A22 tolerance in *sucC* and *mdh* mutants work through different mechanisms. It will be interesting in the future to perform metabolite analysis on the *sucC* deletion strain to better understand the changes in metabolites. Do mutations in other steps of the TCA cycle increase gluconeogenesis? It would seem that they do not as only 3 of the 8 reactions of the TCA lead to increased A22 tolerance when broken. Metabolite analysis of the *sucC* and *acnB* mutants could also help to unravel the differences in mecillinam resistance. Perhaps the changes in ATP levels in the *acnB* mutant provides a general antibiotic resistance mechanism. However, SucC and Mdh occur after the formation of reducing power in the TCA cycle. While the substrate of Mdh (malate), can be directly converted into gluconeogenesis substrates, succinyl-CoA the substrate of SucC, is more likely to be consumed in the modification of amino acids; arginine degradation, and methionine and lysine biosynthesis. It is possible the levels of these specific amino acids relate to A22 resistance. Unfortunately, we were unable to record PEP or oxaloacetate levels in our metabolomic dataset. This leaves open the possibility that other metabolic changes have occurred that account for the increase in the antibiotic tolerance in the *mdh* mutant.

Previous research has shown that while persister cells are more tolerant of aminoglycoside antibiotics, the addition of different sugars specifically potentiated the effects of gentamycin and other aminoglycosides, resulting in a reduction of persister cell survival after treatment (Allison, Brynildsen et al. 2011). The work presented here suggests that the addition of glucose has the opposite effect on the efficacy of mecillinam and decreases its effectiveness. Additionally, inhibiting ATP production can lead to increases in antibiotic tolerance which can be bypassed through the addition of glucose (Conlon, Rowe et al. 2016, Shan, Brown Gandt et al. 2017, RoweRowe, Wagner et al. 2020). While the addition of glucose may help to treat cells when using aminoglycosides by killing both actively growing cells and persister cells, the opposite is true when using mecillinam, as the addition of glucose actually increases the required dose (MIC).

Furthermore, gluconeogenic metabolism has been shown to promote infection. Colonization by *E. coli* of both cows and mice has been shown to increase during gluconeogenic growth, especially when undergoing competition in the gut (Miranda, Conway et al. 2004, Bertin, Deval et al. 2014). Moreover, a gluconeogenic environment leads to an increase in virulence factor expression of enterohemorrhagic *E. coli* (Njoroge, Nguyen et al. 2012). Therefore, the use of metabolites to potentially help eliminate a subset of cells, such as persister cells, may have the unintended consequences of both increasing colonization and virulence of other cells and diminishing the efficacy of other antibiotics.

The properties that make MreB an attractive drug target — conservation across many pathogens and essentiality — are true for other members of the MreB complex, including RodA and RodZ. Our results suggest that drugs that target these proteins would be less effective if gluconeogenesis is activated. As new antibiotics are developed against novel targets, these results show it is important to think about the broad function of the target protein because the efficacy of a drug may be affected in seemingly unknown ways, as we have shown that antibiotic tolerance against MreB-targeting drugs can be increased through a mutation in the TCA cycle gene *mdh*.

## Material and Methods

### Bacterial Growth

Bacteria were grown using standard laboratory conditions. Cultures were grown overnight in LB medium (10 g/L NaCl, 10 g/L tryptone, 5 g/L yeast extract), subcultured 1:1000, and grown to exponential phase (O.D._600_ 0.3-0.6) at 37°C in a shaking incubator.

### Bacterial Strains

All gene deletions in MG1655 were made by P1 transduction from the Keio collection. Transductants were selected on kanamycin (30 μg/mL) and confirmed by PCR. To produce double mutants, pCP20 (Table S1) was transformed into strain 1 in order to remove the kanamycin resistance cassette. P1 transduction was then used to move the second mutation into the strain. See Table S1 for a list of strains used in this study.

*mdh* was cloned into pBad33 by PCR and digestion-ligation at the *EcoRI* and *HindIII* sites. The ligation mixture was used to transform S17 cells and sequence verified before being transformed into MG1655.

### MIC assay

Optical densities of all cultures used in the minimum inhibitory concentration assays (MIC) were checked at 600 nm (OD_600_) with a Thermo Scientific Biomate-3S. Overnight cultures were grown in 2mL test tubes in LB medium in a shaking incubator and normalized to have the same O.D._600_ of the slowest growing culture. 1:100 dilutions were made into 96-well plates filled with 100μl of LB medium plus indicated antibiotics and/or specified sugars or tricarboxylic acids. Two-fold dilutions of each antibiotic were made. Each plate had one row left uninoculated for a blank and one row drug free as a growth control. The O.D. from the drug free wells were halved and used to determine the MIC values as the concentration of drug that caused half the growth as that in the no drug control. Experiments were performed at minimum in triplicate.

### Microscopy

For all imaging, cells were grown at 37°C in indicated medium. Imaging was done on 1% M63-agarose pads at room temperature. Images were collected on a Nikon Ni-E epifluorescent microscope equipped with a 100X/1.45 NA objective (Nikon), Zyla 4.2 plus cooled sCMOS camera (Andor), and NIS Elements software (Nikon).

IDD was calculated using the matlab software Morphometrics (Ursell, Lee et al. 2017), and custom software as previously described (Morgenstein, Bratton et al. 2015, Bratton, Shaevitz et al. 2018). Briefly, a centerline is drawn through each cell and the diameter is measured across the cell body. The standard deviation of these center lines is divided by the mean width of the cell to calculate a coefficient of variation of intracellular diameter deviation. Only non-dividing cells were used for analysis.

Total cell fluorescence was calculated using custom Matlab software. Cell contours were made using Morphometrics to determine the cell boundaries. The total fluorescence within this contour was divided by the amount of pixels to determine the cell brightness

### Metabolomics

#### Collection

Cells were grown overnight shaking at 37°C in LB medium and then subcultured 1:1000 into a flask with 10 ml of fresh LB and grown shaking at 37°C for ~4 hours until in exponential phase (O.D._600_ 0.3-0.4). Cells were passed through a 0.2 μm filter via vacuum filtration. The filter was placed in a glass dish with cold acetonitrile:methanol:water (40:40:20) + 0.5% formic acid and placed at −20°C for 15 minutes to quench metabolism in the cells. This quenching solution was used to wash the cells off of the filter before the solution was neutralized with 1.9M ammonium bicarbonate. The cells were centrifuged at 15,000 rpm for 5 minutes to remove debris. Supernatants were collected for mass spectrometry.

#### Analysis

Supernatants were analyzed via the methods of Su et al. (Su, Chiles et al. 2020). Specificity was achieved through a combination of chromatographic separations followed by high-resolution MS. This method allows for the identification of ~300 water soluble metabolites. Covariant ion analysis (COVINA) was used to identify peaks. Cells were run in triplicate and the effect of *mdh* deletion on metabolites was only determined for metabolites that were identified in all three replicates and showed a consistent ratio between WT: *mdh* cells.

## 3 Author Contributions

BB performed and designed experiments and interpreted data. AG performed experiments. RMM designed experiments, interpreted data, and wrote the manuscript.

## 5 Acknowledgments

The authors would like to thank Dr. Elizabeth Ohneck, and Dr. Tyrrell Conway for editing of this manuscript. They would also like to thank the Rutgers Cancer Institute of New Jersey for help with metabolomics.

## 1 Data Availability Statement

The datasets [GENERATED/ANALYZED] for this study can be found in the [NAME OF REPOSITORY] [LINK]. Please see the [Data Availability section of the Author guidelines](https://www.frontiersin.org/about/author-guidelines#AvailabilityofData) for more details.

